# Functional knockout of forebrain protein 14-3-3 disrupts conditioned taste aversion learning

**DOI:** 10.1101/004663

**Authors:** Adam Kimbrough, Yuying Wu, Yi Zhou, Thomas A. Houpt

## Abstract

Protein 14-3-3 isoforms are key to many cellular processes and are ubiquitous throughout the brain. 14-3-3 is a regulator of ser/thr phospho-signaling by binding and sequestering phosphorylated substrates including kinases, histone deactylases, and transcription factors. The role of protein 14-3-3 in conditioned taste aversion learning (CTA) has not previously been examined. We parameterized CTA learning in difopein-YFP transgenic mice, which have widespread by expression of the artificial peptide difopein in the forebrain, including the basolateral amygdala and insular cortex, resulting in functional knock-out (FKO) of all 14-3-3 isoforms. We found that a single pairing of saccharin or NaCl (CS) and LiCl injection (US) was not sufficient to induce CTA in FKO mice. Multiples pairings of CS and US did lead to CTA acquisition in the FKO mice; however, the CTA rapidly extinguished within 30 minutes to 24 hours after acquisition. Additionally, we found that 14-3-3 FKO mice have an attenuated visceral neuraxis response to LiCl as measured by c-Fos induction. The deficit in FKO was not due to an inability to discriminate or avoid tastants, because they showed normal unconditioned taste preferences for both palatable (saccharin, maltodextrin, low concentration NaCl) and unpalatable tastants (quinine, HCl, and high concentration NaCl) and they were able to reduce intake of a maltodextrin solution adulterated with quinine. The FKO did not have a global deficit in ingestive learning, because they were able to form a conditioned flavor-nutrient preference. Thus, FKO of forebrain 14-3-3 appears to disrupt CTA learning leading to forgetting, rapid extinction, or failure to reconsolidate. This further implicates ser/thr phospho-signaling pathways in the regulation of long-term CTA learning.

## Introduction

Conditioned taste aversion (CTA) is a form of associative learning in which a palatable taste (conditioned stimulus; CS) is paired with a toxic unconditioned stimulus (US) resulting in a reduced preference for the CS. A robust CTA can be formed with one pairing of the CS and US (Garcia and Kimeldorf 1957). Multiple pairings of a CS and US can lead to a stronger CTA (Riley and Mastropaolo 1989).

Several intracellular signaling pathways contribute to CTA learning, including the cyclic-AMP (cAMP)-Protein Kinase A (PKA)-cAMP response element-binding protein (CREB) pathway mediated by ser/thr phospho substrates. When PKA in the amygdala was inhibited long-term, but not short-term CTA memory was attenuated (Koh et al. 2002). Conversely, blocking protein phosphatases PP1/2A in the amygdala via okadaic acid (Oberbeck et al. 2010) or transgenic knockdown of calcineurin (Baumgartel et al. 2008) results in strengthened CTA learning. Within the nucleus, CREB, a substrate of PKA, mediates gene transcription critical for learning. Disruption of CREB in the amygdala leads to attenuated CTA learning (Lamprecht et al. 1997; Josselyn et al. 2004). CREB activation is also seen in the insular cortex (IC) and amygdala during CTA learning as levels of phospoCREB (pCREB) are elevated (Desmedt et al. 2003), and in induction of CREB-regulated immediate early genes such as c-Fos (Koh and Bernstein 2005; Bernstein and Koh 2007; Barot et al. 2008; Kwon et al. 2008; Kwon and Houpt 2010) and ICER (Spencer and Houpt 2001).

14-3-3 proteins are regulatory proteins that form homo-and heterodimers and bind to phosphoserine/threonine sites on target proteins (Jin et al. 2004; Obsilova et al. 2008; Steinacker et al. 2011). 14-3-3 proteins consist of seven isoforms in mammals, which are highly expressed in the brain among other tissues (Watanabe et al. 1991; Watanabe et al. 1993; Watanabe et al. 1994; Steinacker et al. 2011; Lesscher et al. 2012). 14-3-3 can act to sequester bound proteins from other binding partners or restrict subcellular localization to the cytoplasm. 14-3-3 interacts with proteins important in numerous cellular processes such as cellular communication and signal transduction, energy and metabolism, protein synthesis processing and fate, nucleic acid synthesis and processing, cellular organization, and storage and transportation (Jin et al. 2004). 14-3-3 disfunction has been implicated in various neural diseases such as Parkinson’s disease, amyotrophic lateral sclerosis (ALS) Alzheimer’s disease, schizophrenia, and acute neurodegeneration among others (Foote and Zhou 2012). Protein 14-3-3 required for aggresome targeting of misfolded proteins, such that deficiency in 14-3-3 binding may contribute to neurodegenerative diseases (Xu et al. 2013; Jia et al. 2014).

Additionally, 14-3-3 is involved in various aspects of plasticity. Mutation of the *Drosophila* gene *leonardo (leo)*, which encodes one (ζ) of two 14-3-3 isoforms in flies, causes deficits in olfactory learning (Skoulakis and Davis 1996) and disrupts short-term plasticity at the neuromuscular junction (Broadie et al. 1997). Knockdown of 14-3-3 proteins also causes deficits in forms of plasticity in mice, such as hippocampal-dependent plasticity and spatial learning (Nelson et al. 2004; Cheah et al. 2012; Patil et al. 2012; Sun et al. 2013; Qiao et al. 2014), addiction models (Kao et al. 2011; Lesscher et al. 2012) and long-term potentiation (Qiao et al. 2014). Additionally, NMDA receptors, key in many types of plasticity, also interact with protein 14-3-3. Functional knockout of protein 14-3-3 transgenic causes reductions in NMDA receptor levels in the mouse hippocampus (Qiao et al. 2014), while genetic reduction of NMDA receptor levels is accompanied by synapse-specific reductions in 14-3-3ε in the mouse striatum (Ramsey et al. 2011).

Because 14-3-3 sequesters ser/thr phospho-substrates, it is a key regulator of these phospho-signaling pathways. Some ser/thr phospho-substrates involved in CTA learning also have demonstrated interactions with protein 14-3-3, such as MEKK (Kwon and Houpt 2012; Saha et al. 2012) and histone deacetylases (HDACs) (Grozinger and Schreiber 2000; Nishino et al. 2008; Kwon and Houpt 2010; Morris et al. 2013).

To determine if 14-3-3 is required for CTA learning, we tested transgenic mice that express a neuronal specific YFP-fused difopein peptide (Foote and Zhou 2012; Sun et al. 2013; Qiao et al. 2014). Difopein (dimeric 14-3-3 peptide inhibitor or R18) is an artificial peptide sequence with high-affinity binding to all 14-3-3 isoforms (Wang et al. 1999), resulting in functional protein 14-3-3 knockout (FKO) by competitive antagonism in brain regions expressing difopein. A major advantage of difopein-expressing FKO mice have advantages over other methods of 14-3-3 knockdown such as RNAi or viral knockdown is that difopein targets all 7 isoforms of protein 14-3-3. From several lines of FKO mice were developed and we selected a line showing high levels of difopein-YFP expression in the basolateral amygdala (BLA) and IC, areas of relevance for CTA learning.

We used two-bottle preference tests as a measure of CTA learning. Repeated two-bottle preference testing was used to measure the rate of extinction of the learned CTA. CTA strength and persistence were measured in FKO mice compared to wildtype (Wt) mice after pairing palatable tastants with different doses of LiCl and after multiple CS-LiCl pairings. The neuronal response to the US was examined with c-Fos immunohistochemistry in the visceral neuraxis after LiCl injection. Basic responses to palatable and aversive tastants, and the ability of FKO mice to acquire a different form of ingestive learning, conditioned flavor preference, were also assessed.

## Results

### Difopein-YFP distribution

We examined the brains of difopein-YFP mice from line 132 to determine areas of transgene expression. Difopein-YFP was present in both cell bodies and fibers. In areas of relevance to CTA learning, we did not find high levels of difopein-YFP expression in the hindbrain, and noted the absence of expression from the NTS, AP and solitary tract (and hence vagal afferent fibers as well) (see Figure 1A) and the PBN (see Figure 1B). In the forebrain, we found difopein-YFP expressed at very high levels in the BLA (see Figure 1C), and in apparent layer V pyramidal cells of the cortex including IC (see Figure 1D).

**Figure 1.**
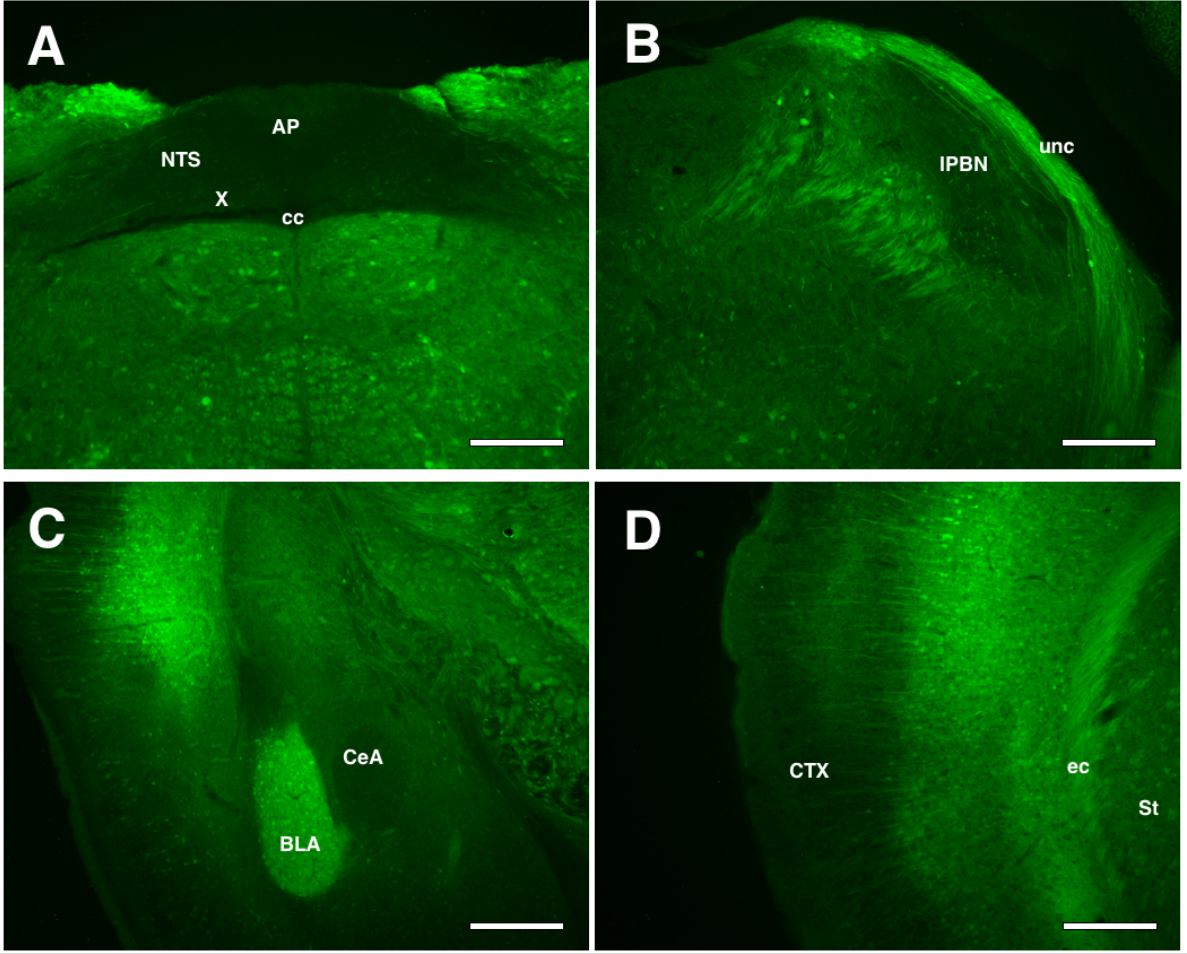
A-D. Fluorescent images of difopein-YFP expression transgenic mice. Difopein-YFP is absent from the NTS, AP and vagal afferents (A) and cell bodies of the PBN (B). There is strong difopein-YFP expression in the BLA (C) and IC (D). Scale bar in all images: 200 microns. (AP) area postrema; (NTS) nucleus of the solitary tract; (cc) central canal; (X) dorsal motor nuclei; (unc) uncinate fasciculus; (lPBN) lateral parabrachial nucleus; (BLA) basolateral amygdala; (CeA) central nucleus of the amygdala; (CTX) cortex, (ec) external capsule; (St) Striatum.

### Experiment 1. Failure to show CTA after a low dose of LiCl

We tested whether 14-3-3 FKO mice had disrupted CTA learning after a single pairing of saccharin and a low dose of LiCl. We paired 10-min access to 0.125% saccharin with 20 ml/kg LiCl or NaCl injection in Wt and FKO mice. The average saccharin intake on conditioning day was 1.8 ± 0.1g. Two-way ANOVA of conditioning day saccharin intake showed no effect of genotype and no interaction, however an effect of drug treatment was seen (F(1,26)=5.46, p < .05). Post-hoc analysis with a Tukey’s HSD showed Wt LiCl mice had an increased intake during conditioning compared to FKO NaCl mice (2.1 ± 0.1 g vs. 1.7 ± 0.1g, p < 0.05)

Two-way repeated measures ANOVA of saccharin preference across testing days showed an interaction of group and days (F(9,75) =4.83, p < .05), such that Wt LiCl treated mice formed a weak CTA but FKO LiCl treated mice did not (see Figure 2A). The CTA of Wt LiCl mice extinguished within a day.

**Figure 2.**
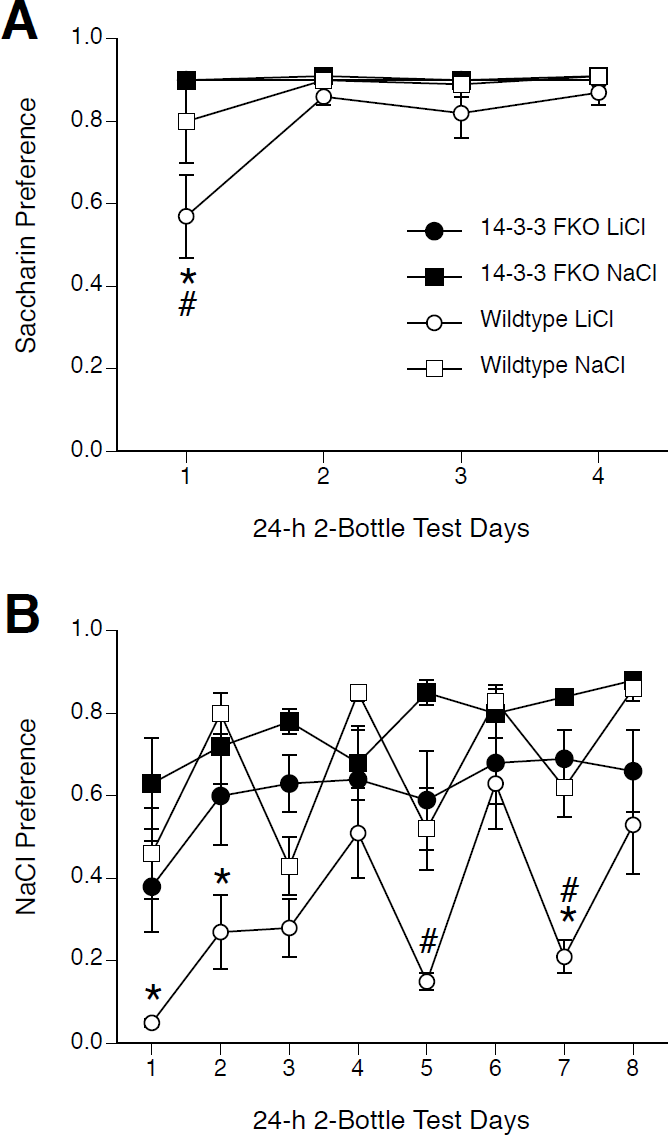
A. Saccharin preference during 24-h 2-bottle extinction testing over 4 days. All mice showed a high preference for saccharin, however Wt-LiCl mice (white circles) showed a significantly lower preference for saccharin on day 1 compared to Wt-NaCl (white squares) mice and FKO-LiCl mice (black circles). The reduced preference seen on day 1 was extinguished by day 2 and no differences were seen for the remainder of 24-h 2-bottle testing. B. NaCl preference during 24-h 2-bottle extinction testing over 8 days. Wt and FKO mice injected with NaCl showed a moderate preference for NaCl that is maintained throughout testing. Wt-LiCl mice (white circles) showed a significantly lower preference for NaCl on days 1, 2 and 7 compared to Wt-NaCl mice (white squares), but FKO-LiCl mice (black circles) failed to show a CTA during testing. * p < .05-Wt-LiCl vs. Wt-NaCl, # p < .05 Wt-LiCl vs. FKO-LiCl.

### Experiment 2. Failure to show CTA after a high dose of LiCl

The 14-3-3 FKO mice in experiment 1 failed to acquire a CTA to saccharin paired with a low dose of LiCl (0.15M, 20 ml/kg). Mice were also tested with 8% maltodextrin paired with LiCl (0.15M, 40 ml/kg) but no aversion was formed in either Wt or FKO mice (data not shown). Therefore, we tested if the 14-3-3 FKO mice could learn a CTA to a less-preferred taste (75mM NaCl) paired with a stronger dose of LiCl (0.15M, 40 ml/kg).

We paired 75 mM NaCl with 40 ml/kg LiCl or NaCl injections in Wt and FKO mice. The average conditioning day NaCl intake for all mice was 2.1 ± 0.1g. Two-way ANOVA of conditioning day NaCl intake showed no effect of genotype, treatment or any interaction.

Two-way repeated measures ANOVA of NaCl preference during 2-bottle preference testing showed an interaction (F(21,196)=2.16, p < .05) (see Figure 2B). Wt mice formed a significant CTA for 75 mM NaCl after pairing with 40 ml/kg LiCl. The CTA extinguished after 2 days in Wt mice. However, FKO mice failed to show a significant CTA on any day. Thus, under conditions in which Wt mice formed a CTA, FKO mice did not acquire a CTA.

### Experiment 3. Acqusition of CTA only after multiple conditioning trials

14-3-3 FKO mice failed to express a CTA after a single pairing of saccharin or NaCl with LiCl injection (experiments 1 and 2). Even in Wt mice, however, the CTA induced by single pairings was transient and rapidly extinguished. Therefore, we tested CTA acquisition and expression in 14-3-3 FKO mice after multiple pairings of saccharin and LiCl (see Table 1).

**Table 1.**
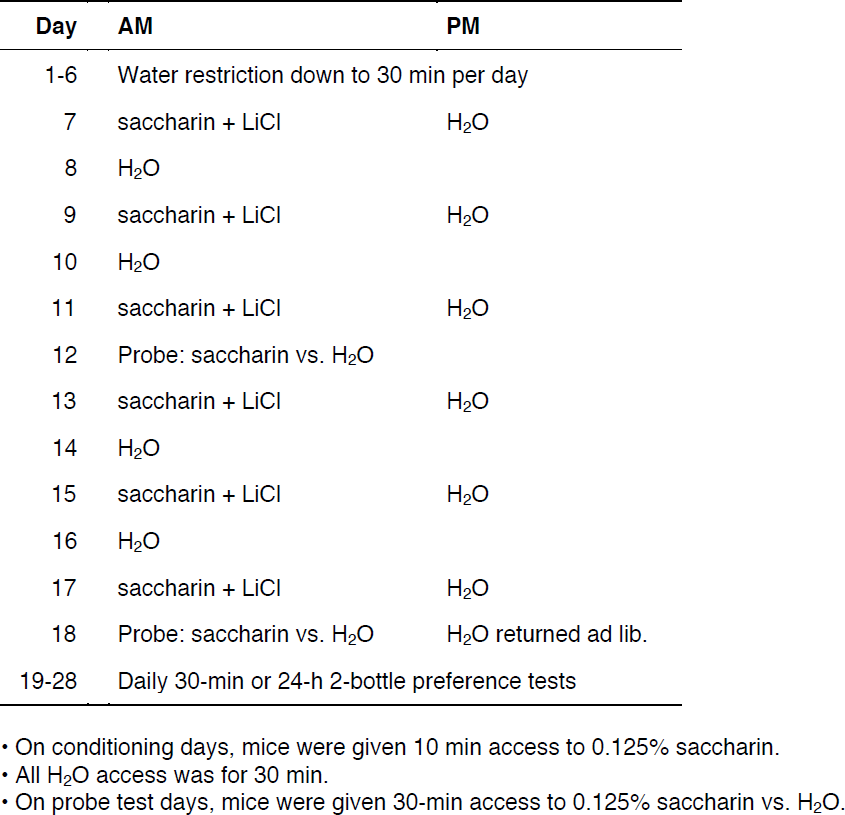
Outline of procedure in Experiment 3.

We performed repeated pairings of saccharin and 20 ml/kg LiCl or NaCl injections on both Wt and FKO mice. Across the 6 pairings, average saccharin intake declined in LiCl-treated mice from day 1 (2.2 ± 0.1g vs. 2.0 ± 0.1 g in NaCl-treated mice, n.s.) until it was significantly lower on day 6 (1.5 ± 0.1 g vs. 2.3 ±. 0.1 g in NaCl-treated mice, p <0.05) No differences between genotypes were seen during conditioning.

Two-way repeated measures ANOVA of saccharin preference on the two 30-min 2-bottle probes revealed an effect of group (F(3,52)=23.23, p < .05), and day (F(1,52)=10.79, p < .05), but no interaction (see Figure 3A). All mice treated with LiCl acquired a CTA after 3 and 6 pairings of saccharin and LiCl, which persisted through both probe test days. Thus, FKO mice were able to form a CTA after multiple pairings of saccharin and 20 ml/kg LiCl.

**Figure 3.**
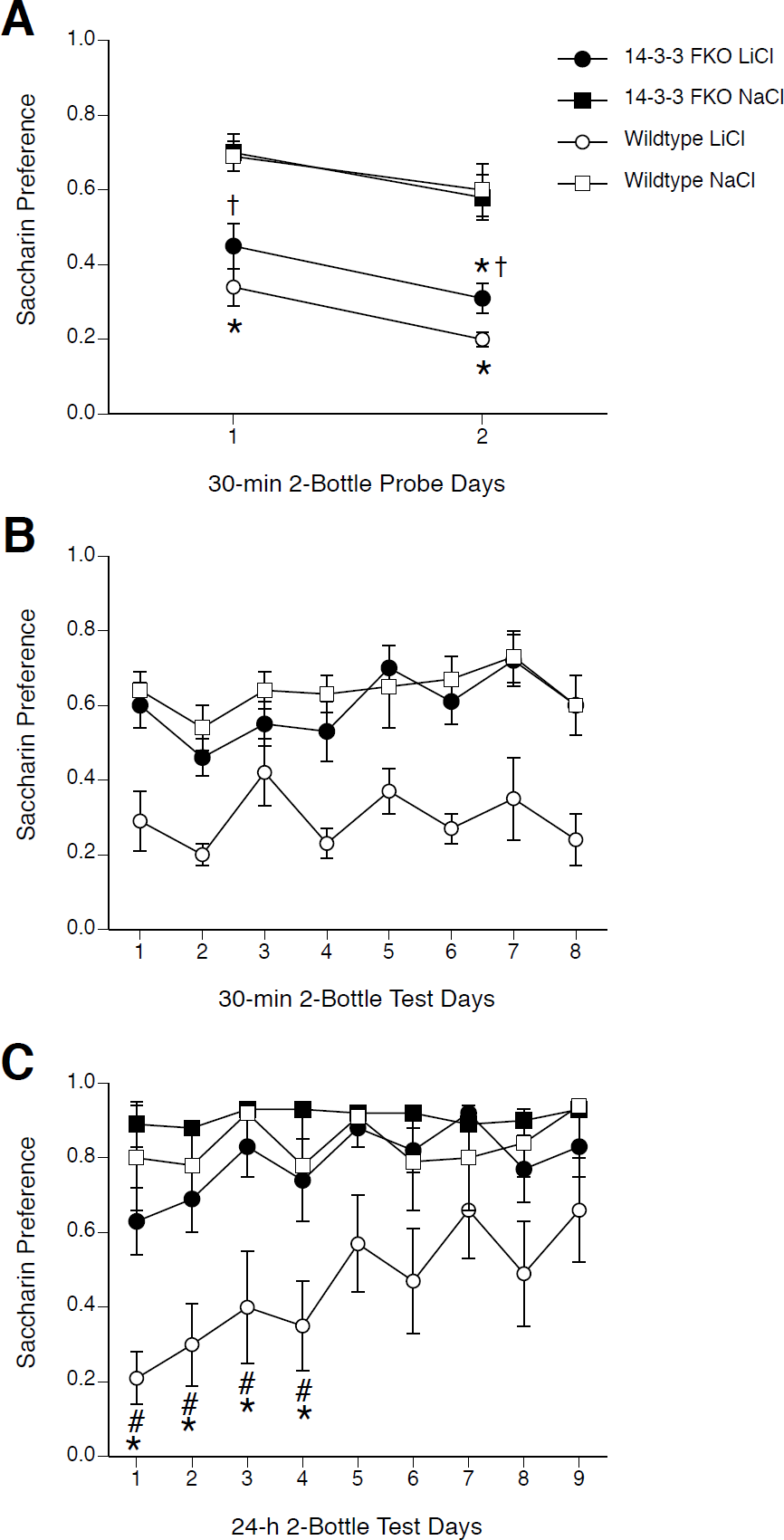
A. Saccharin preference during 30-min 2-bottle probe test days after 3 and 6 pairings of saccharin and 20 ml/kg LiCl. Wt and FKO mice injected with NaCl showed a preference for saccharin on both probe days 1 and 2. Both Wt-LiCl (white circles) and FKO-LiCl (black circles) mice had a significantly reduced preference for saccharin compared to Wt-NaCl (white squares) or FKO-NaCl (black squares) mice on both probe days 1 and 2. B. Saccharin preference during 30-min 2-bottle extinction testing over 8 days. FKO-LiCl (black circles) and NaCl-injected (white squares) mice showed a high preference for saccharin. Wt-LiCl mice (white circles) had a low preference for saccharin throughout extinction testing, although this preference was not significantly different from NaCl-injected or FKO-LiCl mice. C. Saccharin preference during 24-h 2-bottle extinction testing over 9 days. FKO-LiCl (black circles), FKO-NaCl (black squares), and Wt-NaCl (white squares) mice showed a high preference for saccharin. Wt-LiCl mice (white circles) had a significantly lower preference for saccharin compared to Wt-NaCl and FKO-LiCl mice on days 1 through 4. Thus, the Wt-LiCl mice formed a CTA that persisted for 4 days and extinguished by day 5, whereas FKO-LiCl showed no CTA during extinction testing. * p < .05-Wt-LiCl or FKO-LiCl vs. Wt-NaCl, † p < .05-FKO-LiCl vs. FKO-NaCl.

#### Experiment 3a. Rapid extinction of CTA across 30-min sessions

Across 8 days of 30-min 2-bottle preference testing, two-way repeated measures ANOVA of saccharin preference showed an effect of group (F(2,21)=37.57, p<.05), and day (F(7,47)=2.42, p<.05), but no interaction (see Figure 3B).

Comparing group preferences averaged across all 8 extinction days showed a significant effect of group (F(2,21)=36.76, p <0.05), with Wt LiCl showing a significantly lower preference than NaCl-treated mice and FKO LiCl mice (which were not different from NaCl-treated mice).

Thus, both Wt and FKO mice formed a CTA after multiple pairings of saccharin and 20 ml/kg LiCl. However, the CTA was extinguished in the FKO mice by the first 30-min test following probe testing.

#### Experiment 3b. Rapid extinction of CTA across 24-h sessions

Across 8 days of 24-h 2-bottle preference testing, two-way repeated measures ANOVA of saccharin preference showed an effect of group (F(3,28)=10.04, p < .05), and an effect of day (F(8,224)=5.10, p < .05) but no interaction (see Figure 3C). Tukey’s HSD post-hoc analysis showed Wt LiCl mice had a significantly lower preference on days 1 through 4 compared to Wt NaCl mice and FKO LiCl mice. (Analysis of saccharin intake showed an interaction of group and days (F(24,224)=2.5, p < .05), results not shown.)

Thus, both FKO and Wt mice acquired a CTA (as seen in probe testing), but the CTA persisted in Wt mice while rapidly extinguishing in FKO mice. Thus, 14-3-3 FKO mice may have not only a deficit in acquisition of a CTA, but also deficits in long-term retention or extinction.

### Experiment 4. Unconditioned taste aversions and preferences

#### Experiment 4a. Normal unconditioned taste preferences

Mice were tested for unconditioned taste aversions and preferences to determine if 14-3-3 FKO mice were able to reduce intake of unconditioned tastants that are normally rejected by Wt mice and increase intake of tastants that are preferred by Wt mice. To examine unconditioned taste preferences of 14-3-3 FKO mice, we measured intake of water vs. various tastants in 48-h 2-bottle preference tests.

Like Wt mice, FKO mice showed a preference for 0.125% saccharin, 8% maltodextrin, and 75 mM NaCl over water, and an aversion to 600 mM NaCl, 30 *µ*M quinine sulfate, and 10 mM HCl (see Figure 4A). There was no difference between Wt and FKO preferences except that FKO mice showed a small but significant decrease in preference for 10 mM HCl. Thus, FKO mice displayed innate aversions or preferences similar to those of Wt mice for various tastants.

**Figure 4.**
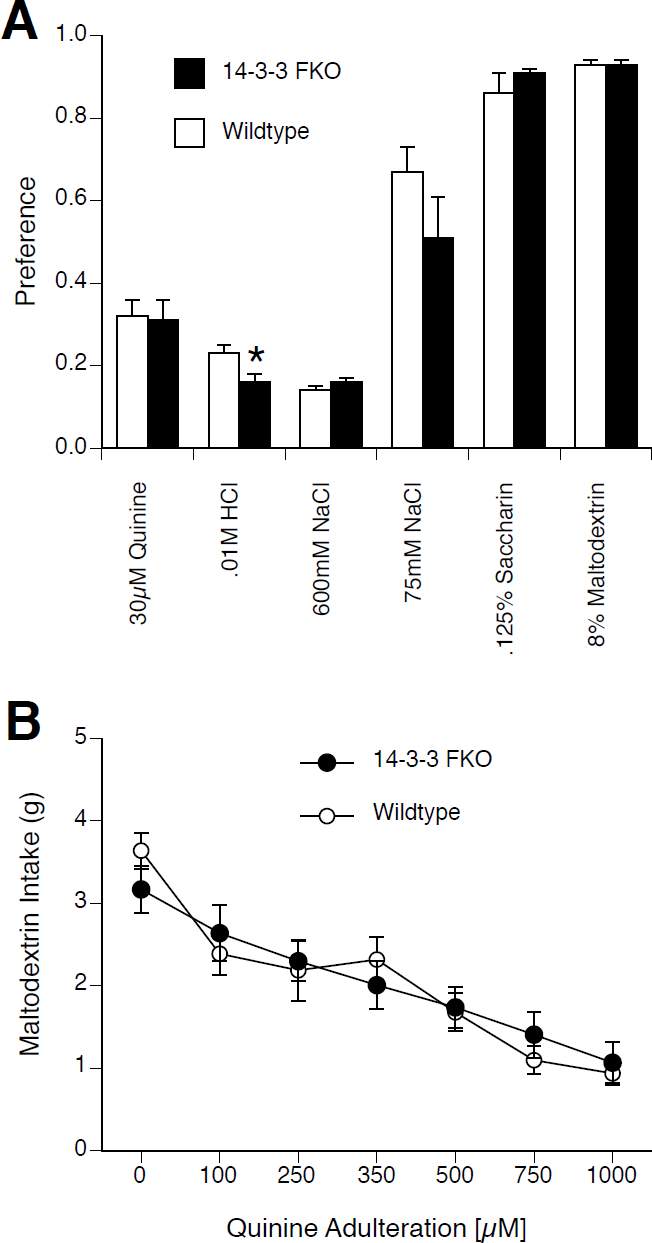
A. 48-h preference for unconditioned tastants. Both Wt (white bars) and FKO mice (black bars) had high preferences for 75 mM NaCl, 0.125% saccharin, and 8% maltodextrin. Both Wt and FKO mice had low preferences for 30 *µ*M quinine sulfate, 0.01M HCl, and 600 mM NaCl. FKO mice showed a significantly lower preference for 0.01M HCl than Wt mice. B. 2-h single bottle 8% Maltodextrin intake with increasing concentrations of quinine adulteration. With no quinine present both Wt (white circles) and FKO mice (black circles) had a high 8% maltodextrin intake. Wt mice significantly reduced intake of 8% maltodextrin with addition of quinine at all concentrations. FKO mice significantly reduced intake of 8% maltodextrin with addition of quinine at concentrations of 250 *µ*M and higher. * p < .05 Wt vs FKO.

#### Experiment 4b. Avoidance of quinine-adulterated solution

We also measured intake by FKO and Wt mice of palatable maltodextrin adulterated with increasing concentrations of quinine (Lesscher et al. 2012). Two-way repeated measure ANOVA revealed an interaction of genotype and quinine concentration (F(6,180)=2.41, p < .05) (see Figure 4B). Post-hoc analysis with Tukey’s HSD at each quinine concentration showed no differences between FKO and Wt mice. For Wt mice, quinine at all concentrations significantly reduced 8% maltodextrin intake compared to baseline. For FKO mice, 250 mM quinine (but not 100 mM quinine) and higher reduced 8% maltodextrin intake compared to baseline. Thus, both Wt and FKO mice were able to lower their intake of a palatable solution in response to the presence of an increasing concentration of a bitter tastant (i.e. without conditioning).

### Experiment 5. Normal conditioned flavor preference learning

To determine if another form of taste-guided learning showed deficits in 14-3-3 FKO mice similar to those seen in CTA learning, we tested 14-3-3 FKO mice in a conditioned flavor preference (CFP) learning paradigm.

During 48-h access to CS+/8% glucose solutions, mice drank CS+ almost exclusively (46.8 ± 0.6 g CS+ vs. 3.8 ± 0.3 g water). Subsequently, on the first day of 2-bottle testing both FKO and Wt mice showed a high preference for the CS+ without glucose vs. CS-, indicating the ability to learn a CFP. Two-way repeated measures ANOVA of CS+ preference during 2-bottle testing showed an effect of group (F(1,27)=6.29, p < .05), an effect of day (F(5,135)=4.03, p < .05), and an interaction (F(5,135)=2.46, p < .05) (see Figure 5). There was no difference between Wt and FKO day 1 preferences compared to day 6 preferences for the same group (i.e. Wt to Wt and FKO to FKO). This indicates that both FKO and Wt CS+ preferences did not extinguish over the 6 days of 2-bottle testing. Furthermore, the only significant difference between FKO and Wt mice occurred on day 2; when analyzed without data from day 2 data, there was no significant effects of genotype or day on preference.

**Figure 5.**
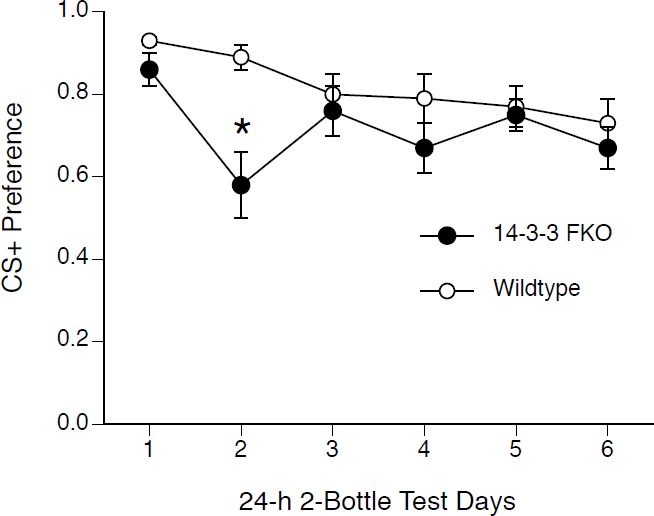
CS+ preference during 24-h 2-bottle extinction testing after 48-h CFP conditioning. Both Wt (white circles) and FKO mice (black circles) showed a high CS+ preference on day 1 which was not significantly different from CS+ preference on day 6. FKO mice showed a significantly lower CS+ preference on day 2 compared to Wt mice. * p < .05 Wt vs FKO.

Two-way repeated measures ANOVA of CS+ intake during 2-bottle testing showed an interaction of genotype and day (F(5,135)=5.25, p < .05) (data not shown). Wt and FKO mice both showed a decreased CS+ intake on day 6 compared to day 1 for the same group (i.e. Wt to Wt and FKO to FKO). Thus, although they did not show extinction in preference, both Wt and FKO mice did show extinction of CS+ intake over the course of 2-bottle testing.

### Experiment 6. Attenuated c-Fos induction after LiCl injection

To access neural processing of the US and activation of the visceral neuraxis, we examined immunohistochemistry of c-Fos after LiCl injection. We examined c-Fos induction in Wt and FKO mice 1 h after LiCl injection (see Figure 6). By two-way ANOVA, significant effects of either drug treatment or genotype were found on LiCl-induced c-Fos expression in the NTS (interaction: F(1,16)=5.7 p<.05), CeA (drug treatment: F(1,15)=26.73, p<.05), SON (drug treatment: F(1,14)=26.0, p<.05, genotype: F(1,14)=5.1, p <0.05), and PVN (drug treatment: F(1,17)=11.1, p<.05), but not in the PBN. In all brain regions except the PBN, LiCl induced significantly more c-Fos compared to NaCl injection in Wt mice. In FKO mice, LiCl induced intermediate levels of c-Fos in the NTS and PVN, and equivalent c-Fos in the CeA, compared to Wt mice. Thus, there appears to be a deficit in processing of the US by the visceral neuraxis in FKO mice.

**Figure 6.**
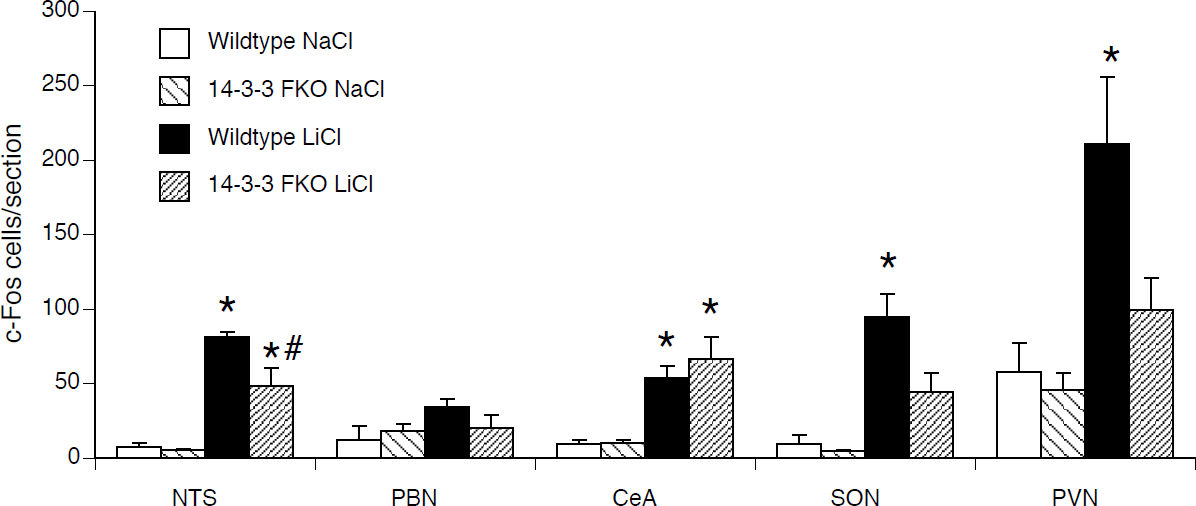
c-Fos counts from Wt and FKO mice in the visceral neuraxis 1 h after LiCl injection. LiCl injection led to increased c-Fos of Wt mice in all brain regions except the PBN. In FKO mice c-Fos was not significantly increased by LiCl injection in the SON, PVN, and PBN. (white) Wt NaCl; (light hatch) FKO NaCl; (black) Wt LiCl; (dense hatch) FKO LiCl. * p < .05-Wt-LiCl or FKO-LiCl vs. Wt-NaCl, † p < .05-FKO-LiCl vs. FKO-NaCl, # p < .05 Wt-LiCl vs. FKO-LiCl.

## Discussion

14-3-3 FKO mice showed a deficit in acquisition or expression of a CTA after a single long-delay pairing. As an initial screen of CTA learning, we used a single long-delay pairing of saccharin and low-dose LiCl, under conditions which demonstrate the prototypical features of CTA and induce a minimal but reliable CTA in Wt mice. FKO mice failed to reduce saccharin preference after a single long-delay pairing, as measured in 24-h 2-bottle preference tests. Strengthening the US to 40 ml/kg LiCl in a single pairing also did not lead to learning.

If the conditioning was strong enough, however, the FKO mice were able to acquire a transient CTA. We tested FKO mice with 6 pairings of the US and CS with a probe test for strength of CTA after 3 pairings and 6 pairings. Both Wt and FKO mice showed a CTA at 24 h after both the 3^rd^ or 6^th^ pairing.

Even after multiple pairings, however, FKO mice have a deficit in CTA learning that appears to be either rapid extinction or forgetting of a learned CTA, or failure to reconsolidate a CTA memory after an expression test. After CTA acquisition induced by multiple pairings the CTA persisted in Wt mice for several days of 30-min or 24 h 2-bottle testing. FKO mice showed a CTA in the 30-min probe test prior to extended extinction testing, however in both 30-min and 24-h 2-bottle tests the CTA was gone on second day of extinction. So, the CTA acquired by the FKO is not expressed beyond a single 30-min 2-bottle session, and is either extinguished or forgotten during the next 24 h.

Given the potential contribution of 14-3-3 to multiple behaviors the basic ingestive responses of the FKO mice were examined. The CTA deficit found in FKO mice was not accompanied by a failure to detect and respond to tastants per se, nor was it due to an inability to reject a tastant. In 2-bottle tests of aversive and palatable solutions Wt and FKO mice showed similar unconditioned taste preferences.

Furthermore, the deficit in CTA learning of FKO mice was not due to the inability of FKO mice to switch from a preference to an aversion of a previously palatable tastant. FKO mice were able to reject a palatable solution that was made increasingly bitter (although others have reported that RNAi knockdown of 14-3-3ζ in the CeA reduced rejection of quinine adulterated ethanol solutions (Lesscher et al. 2012).

We also found that the CTA deficit is not due to a generalized inability to acquire and express conditioned ingestive behaviors. To determine if the deficits seen in CTA extinction extended to other taste-related learning paradigms, we tested mice with CFP learning. Wt and FKO mice both displayed a learned preference for the CS+ flavor of kool-aid paired with glucose which persisted for several days without extinction.

14-3-3 FKO mice, however, have an attenuated visceral neuraxis response to the US. Following LiCl injection, Wt mice showed high c-Fos levels in all brain regions examined as expected. 14-3-3 FKO mice showed similar levels of c-Fos as Wt mice in some brain regions. However, FKO mice showed intermediate c-Fos levels in several brain regions of the visceral neuraxis (NTS, PVN, SON).

Thus, 14-3-3 FKO mice can acquire a CTA, but only after multiple pairings of the CS and US, and the acquired CTA is transient. CTA learning is a multi-faceted process, and multiple deficits may contribute to the impairment found in the FKO mice.

### Cause of the Deficit

The 14-3-3 FKO mice appear to have deficits in extinction, forgetting, or reconsolidation of a learned CTA. We have ruled out several other possible causes of the CTA deficit.

A problem with sensory transduction would impair learning. However, FKO mice preferred palatable tastants and rejected aversive tastants indicating normal gustatory responses to the CS. The attenuated visceral response of FKO mice to LiCl (as measured by c-Fos) indicates a possible problem with detecting the US. Further experiments are required to completely rule out a visceral deficit as a cause for the CTA deficit of FKO mice.

A deficit in associating the CS and US during conditioning would also cause a deficit in subsequent CTA expression. We found FKO mice were able to form a CTA after multiple pairings of the CS and US, but not after only one pairing. This suggests that the FKO mice may have a deficit in associating the CS and US such that only multiple pairings are sufficient for CTA acquisition. Alternatively, single-pairing and multiple-pairing CTA learning could have different mechanisms, such that 14-3-3 FKO mice may have a deficit in single pairing learning, but not multiple pairings.

However, the data points to deficits of either failure to reconsolidate, or rapid forgetting, or rapid extinction. Additional studies will be required to distinguish these possibilities.

### Neurological and Molecular Deficits

In the FKO mice, antagonism of 14-3-3 by expression of the difopein transgene was limited to neurons by the *thy-1* promoter. Furthermore, the specific transgenic line we examined had wide-spread but not ubiquitous difopein-YFP expression in the brain. In particular, although the FKO mice we tested did not express difopein-YFP in many nodes of the visceral and gustatory neuraxes, they did show robust expression in the BLA and IC. We hypothesize that disruption to 14-3-3 in one or both of these regions is responsible for the CTA deficit. Both the BLA and IC are important for CTA learning as demonstrated by many pharmacological, lesion, and molecular studies; indeed, the BLA provides a major and plastic input to the IC that is potentiated by CTA (Escobar and Bermudez-Rattoni 2000; Ferreira et al. 2005). For example, it has been found that lesioning the BLA (Reilly and Bornovalova 2005) or disrupting local CREB gene transcription eliminates CTA learning (Josselyn et al. 2004). Additionally, lesions to the IC attenuate CTA learning, however this may be due to lack of novelty recognition (Koh et al. 2003; Koh and Bernstein 2005; Bernstein and Koh 2007; Roman and Reilly 2007; Roman et al. 2009). Rats with IC lesions acquired CTA slowly regardless of the tastant being novel or familiar, whereas non-lesioned controls acquired a CTA more rapidly with a novel, compared to a familiar, tastant (Roman and Reilly 2007). 14-3-3 knockout in the IC may cause the mice to treat solutions as familiar regardless of experience. Site-specific knockdown of 14-3-3 (or site-specific rescue of 14-3-3 function in the FKO mice) will be required to verify its role in the BLA or IC.

Interestingly, we found attenuated visceral neuraxis responses to LiCl in FKO mice in the SON, PVN, and NTS; although these areas were lacking difopein-YFP expression. This suggests that the attenuated c-Fos response is an effect of the visceral neuraxis network as a whole, such that areas expressing difopein-YFP, e.g. BLA and IC, diminish responsiveness of other parts of the network.

### 14-3-3 in Learning and Memory

Protein 14-3-3 binds ser/thr kinase substrates, and thousands of proteins contain the 14-3-3 binding motif. Over 200 molecules have been shown experimentally to interact with protein 14-3-3 (Jin et al. 2004). However, a few binding partners of protein 14-3-3 have previously been implicated in CTA learning and thus make potential targets for further study.

MAPK can be indirectly modulated, e.g. by MEKK binding, by 14-3-3 proteins in various cellular processes (Wang et al. 1999; Saha et al. 2012; Yan et al. 2013). MAPK inhibition in the CeA has been found to attenuate CTA learning (Kwon and Houpt 2012), and MAPK in the IC is activated during learning of a CTA to a novel taste (Berman and Dudai 2001). MAPK activity in the BLA may also be important for CTA learning, and may be disrupted by 14-3-3 knockout.

HDACs also interact directly with protein 14-3-3 (Grozinger and Schreiber 2000; Nishino et al. 2008), and HDAC inhibition has been found to have varying effects on CTA learning. Systemic inhibition of all HDACs by sodium butyrate during CTA acquisition led to enhanced CTA (Kwon and Houpt 2010). However, specific genetic constitutive knockdown of HDAC 2 diminished CTA learning (Morris et al. 2013). Transducer-of-regulated-CREB-activity (TORC) protein is another binding partner of 14-3-3. TORC is a necessary co-factor for CREB gene transcription (Bittinger et al. 2004), which is sequestered in the cytoplasm in a phosphorylated state by protein 14-3-3 (Screaton et al. 2004). After Ca++ activation, dephosphorylated TORC is released by 14-3-3, translocates to the nucleus, binds to CREB, and facilitates gene transcription (Conkright et al. 2003a; Conkright et al. 2003b; Iourgenko et al. 2003; Bittinger et al. 2004).

NMDA receptors are also critical for CTA learning, and NMDA receptors and protein 14-3-3 can directly or indirectly influence each other’s expression or function. FKO of 14-3-3 decreases NMDA receptors in the hippocampus (Qiao et al. 2014), while knockdown of NMDA receptors reduces 14-3-3ε in the striatum (Ramsey et al. 2011). In CTA learning, NMDA receptors have been shown to be critical and their importance in the IC and BLA has been shown. Increasing NMDA receptor activity by systemic d-cycloserine, an NMDA receptor agonist, (Nunnink et al. 2007; Davenport and Houpt 2009), or by genetic overexpression of the NR2B subunit leads to increased CTA learning (Li et al. 2010). Conversely, antagonism of NMDA receptors attenuates CTA learning. In particular, infusion of CPP or AP5 into the IC can interfere with CTA learning (Ferreira et al. 2005; Rodriguez-Duran and Escobar 2014). In the BLA, infusion of AP5 causes a failure to reconsolidate previously-acquired CTA memory (Garcia-Delatorre et al. 2014). Therefore, disruption of NMDA signaling in the BLA or IC of the FKO might also contribute to their CTA deficeit.

Various protein 14-3-3 isoforms have also been implicated in some specific learning and memory paradigms, although their role has not been thoroughly elucidated in any one system. Hippocampal 14-3-3 γ and ε protein levels have been reported to increase after spatial learning in rats (Nelson et al. 2004; Patil et al. 2012), while 14-3-3ζ deficient mice had attenuated spatial learning ability (Cheah et al. 2012). Conversely, 14-3-3ζ in the nucleus accumbens is downregulated in mice after cocaine-induced conditioned place preference, while overexpression of accumbens 14-3-3ζ diminished expression of the place preference These reported effects of 14-3-3 knockdown are small in magnitude compared to the deficit in CTA learning we saw in the 14-3-3 FKO. The 14-3-3 FKO mice also showed deficits in contextual fear conditioning and passive avoidance, potentially due to functional deficits in hippocampal function (Qiao et al. 2014). These large deficts are perhaps due to the *thy-1* promoter which ensured wide-spread neural expression of the transgene, and because difopein antagonizes all isoforms of 14-3-3 (Kao et al. 2011).

In conclusion, protein 14-3-3 is critical for normal CTA learning as evidenced by the disruption of learning when functionally antagonized. By binding molecules with phospho-ser/thr residues, protein 14-3-3 isoforms play a key role as a gatekeeper in regulating ser/thr and phosphatase signaling (Jin et al. 2004). Molecules bound to protein 14-3-3 may undergo conformational or activation state changes critical for their function (Reinhardt and Yaffe 2013). Protein 14-3-3 also acts as an intermediate step in these signaling pathways by sequestering ser/thr phosphorylated proteins. These proteins can then be released by various protein phosphatases. In CTA learning, several steps in the ser/thr signaling pathway have been examined, primarily involving CREB signaling. PKA, protein phosphatase PP1/2A, calcineurin, and CREB are all involved in CTA learning (Lamprecht et al. 1997; Koh et al. 2002; Josselyn et al. 2004; Baumgartel et al. 2008; Oberbeck et al. 2010). We have now shown the importance of protein 14-3-3 in signaling pathways involved in CTA learning that are regulated by ser/thr phosphorylation.

## Methods

### Animals

Transgenic 14-3-3 FKO mice were generated by expressing YFP-fused difopein using the *thy-1* promoter. Transgenic lines were backcrossed to C57BL/6J mice for at least eight generations. Heterozygous transgenic mice and their WT littermates were identified by PCR genotyping using the following primers: Thy1F (AAGGGGATAAAGAGAGGGGCTGAG) from the *thy-1* sequence, and difopeinR (CTCGCCGGACACGCTGAACTTG) from the difopein sequence. Adult male and female FKO from transgenic line 132 and wildtype (Wt) mice were used for the experiments.

All mice were caged individually and kept on a 12:12 light-dark cycle at 22° C with food and water available ad libitum unless noted otherwise. Experiments were conducted during the light phase of the cycle. All procedures were approved by the Florida State University institutional animal care and use committee.

### Experiment 1. CTA after a low dose of LiCl

The number of mice in each group was: FKO LiCl=8 (4 female, 4 male), FKO NaCl=8 (4 female, 4 male), Wt LiCl=8 (4 female, 4 male), Wt NaCl=6 (4 female, 2 male).

Mice were water restricted for 6 days with access tapered from 3 h/day to 30 min/day. On conditioning day mice were given 10-min access to 0.125% saccharin. 10 min after the end of saccharin access mice were injected with either LiCl or NaCl (0.15M, 20 ml/kg) based on group. Water was returned overnight. The following day 24-h 2-bottle preference tests with water and 0.125% saccharin were begun for 4 days.

### Experiment 2. CTA after a high dose of LiCl

Mice from experiment 1 were used (with 2 additional Wt male mice added so that n=8/group in all groups). Groups were counterbalanced for prior exposure to LiCl as well as balanced for sex.

Mice were water restricted for 6 days with access tapered from 3 h/day to 30 min/day. On conditioning day, mice were given 10-min access to 75mM NaCl. Ten minutes after the end of NaCl access, mice were injected with either LiCl or NaCl (0.15M, 40 ml/kg) based on group. Water was returned overnight ad libitum. The following day, 24-h 2-bottle preference tests with water and 75mM NaCl were begun for 8 days.

### Experiment 3. CTA after multiple conditioning trials

Naïve male and female mice, FKO (n=33) and Wt (n=23) were split into 4 groups (FKO LiCl n=20; FKO NaCl, n=13, Wt LiCl, n=13, Wt NaCl, n=10). Mice were water restricted for 6 days with access tapered from 3 h/day to 30 min/day. On conditioning days mice were then given 10-min access to 0.125% saccharin. 10 min after the end of saccharin access, mice were injected with either LiCl or NaCl (0.15M, 20 ml/kg).

Several hours later, mice were given 30 min of supplemental water. The following day mice received an ‘off’ day and were given 30-min access to water without injection. This cycle of conditioning and ‘off days’ was repeated 3 times. The day following the third conditioning day, mice were given a 30-min 2-bottle preference test with 0.125% saccharin and water instead of an ‘off’ day. The next day the conditioning cycle resumed for 3 more days of conditioning. On the day after the 6th conditioning day a second 30-min 2-bottle preference test was conducted with water and 0.125% saccharin. Water was then returned ad libitum overnight. The next day mice were split into groups and underwent one of 2 schedules of extinction trials with 2-bottle preference tests.

#### Experiment 3a. Extinction across 30-min sessions

For experiment 3a, half of the conditioned mice underwent daily 30-min 2-bottle preference tests. The number of mice (all male) in each group was: FKO LiCl=10, FKO NaCl=4, Wt LiCl=6, Wt NaCl=4. For statistical analysis, the FKO and Wt NaCl-treated mice were combined into a single control group (n=8). Mice were given one 30-min 2-bottle preference test with 0.125% saccharin and water daily for 8 days.

#### Experiment 3b. Extinction across 24-h sessions

For experiment 3b, the other half of the conditioned mice underwent daily 24-h 2-bottle preference tests. The number of mice in each group was: FKO LiCl=10 (5 female, 5 male), FKO NaCl=9 (5 female, 4 male), Wt LiCl=7 (3 female, 4 male), Wt NaCl=6 (3 female, 3 male). Mice were given 24-h 2-bottle preference tests with 0.125% saccharin and water for 9 days.

### Experiment 4 Unconditioned taste aversions and preferences

#### Experiment 4a. Unconditioned taste preferences

Mice from experiments 1 and 2 were used: FKO n=16 (8 female, 8 male) and Wt n=16 (8 female, 8 male)

Mice were given 48-h 2-bottle preference tests with water and one of three tastants (30 micromolar quinine sulfate, 0.01M HCl, 600mM NaCl). Each mouse was tested with each tastant. Preferences were calculated as 48-h tastant intake divided by 48-h total intake (tastant + water).

Additionally, data from the first two 24-h 2-bottle preference tests of NaCl injected mice was used from CTA experiments to provide unconditioned preferences for 0.125% saccharin, 75 mM NaCl and 8% maltodextrin.

#### Experiment 4b. Aversion to quinine-adulterated solution

Mice from experiment 3b were used: FKO n=19 (10 female, 9 male) and Wt n=13 (6 female, 7 male). Mice were given 2-h single bottle access each day, with 2 days of 4% maltodextrin followed by 9 days of 8% maltodextrin. After the 11 days of maltodextrin access, mice were given 2-h single bottle access to 8% maltodextrin adulterated with quinine sulfate. The quinine adulteration lasted for 6 days with quinine concentrations ascending each day (100, 250, 350, 500, 750, and 1000 *µ*M).

### Experiment 5. Conditioned flavor preference learning

Mice from experiments 1, 2, and 4a were used: FKO n=14 (7 female, 7 male) and Wt n=15 (7 female, 8 male).

Two solutions were used: 0.05% Kool-Aid mixed with 8% glucose and 0.05% saccharin (CS+) or a Kool-Aid mixed with saccharin alone (CS-). Grape and cherry flavors were counterbalanced for CS+ and CS-across mice. Mice were given 48-h 2-bottle access to water and the CS+ Kool-Aid flavor. After 48 h, the mice were given 6 days of 2-bottle testing with CS+ and CS-both mixed with only 0.05% saccharin to determine if a preference for CS+ had been learned.

### Experiment 6. c-Fos after LiCl injection

Mice (n=22) from experiments 1,2,4 and 5 were used. Mice were split into 4 groups: FKO LiCl (n=6; 3 male, 3 female), FKO NaCl (n=5; 3 male, 2 female), Wt LiCl (n=6; 3 male, 3 female), and Wt NaCl (n=5; 3 male, 2 female). Mice were injected with either LiCl or NaCl (0.15M, 20 ml/kg) and then perfused 1 h later.

### Tissue Collection

At 1h after LiCl or NaCl injection mice were anesthetized with sodium pentobarbital (104 mg/0.4 ml) and perfused first for 10 min with isotonic saline containing 0.5% sodium azide and 1000 U heparin, and then for 5 min with phosphate-buffered 4% paraformaldehyde. The brains were removed and post-fixed for 24 h, then cryoprotected in 30% sucrose solutions for 1-2 days. The brains were cut at 40 *µ*m at −20°C on a freezing-sliding microtome.

### Immunohistochemistry

Brain sections were washed twice in 0.1 M sodium phosphate-saline (PBS) for 10 min, and then permeabilized in 0.2% Triton-1% BSA-PBS for 30 min. After two PBS-bovine serum albumin (BSA) washes, sections were incubated overnight at 25°C with the appropriate primary antibody.

Sections were incubated in the primary antibody (anti-c-Fos polyclonal antisera: Ab-5, 1:10,000, Calbiochem). Sections were then washed twice with PBS-BSA for 10 min. Sections were then incubated for 1 h with a biotinylated secondary antibody (e.g. goat anti-rabbit antibody (1:200, Vector Laboratories)). Antibody complexes were amplified using the Vectastain ABC Elite kit (Vector Laboratories). Following amplification sections were washed twice with 0.1M phosphate buffer (PB) for 10 min. Immunoreactivity was then visualized by a 5-min reaction with 0.05% 3,3-diaminobenzidine tetrahydrochloride. Sections were then immediately washed twice in 0.1 M PB and mounted on gelatin-coated slides. Slides were coverslipped with Permount.

### Quantification

Cells expressing darkly positive, nuclear staining were quantified using custom software (MindsEye, Thomas A. Houpt). Regions were digitally captured at 40x magnification on a Macintosh computer using an Olympus Provis AX-70 microscope with a Dage-MTI DC-330 CCD camera and Scion LG-3 framegrabber. Counting was restricted to the area delineated by a hand drawn outline. Unilateral cell counts were averaged for sections for each mouse. The individual mean counts for each region were averaged across mice within experimental groups. Unilateral cell counts were averaged for 2 sections of the paraventricular nucleus (PVN; bregma −.70 mm to −.94 mm), 4 sections of the supraoptic nucleus (SON; bregma −.58 to −.94), 4 sections of the central amygdala (CeA; bregma −.70 mm to −1.34 mm), 2 sections of the parabrachial nucleus (PBN; bregma −4.96 mm to −5.52 mm), and 4 sections of the nucleus of the solitary tract (NTS; bregma −7.2 mm to −7.76 mm) (Paxinos and Franklin 2001).

### Statistical analysis

Preference scores and cell counts were analyzed by 2-way analysis of variance (ANOVA) using Statistica software (Statsoft, Tulsa, OK). For CTA extinction (Expt 1-3), maltodextrin preference (Expt 4b), and CFP expression (Expt 5), group (Wt-NaCl, Wt-LiCl, FKO-NaCl, FKO-LiCl) and day of preference testing were the independent factors with CS preference as the repeated dependent measure. Tukey-Kramer HSD pairwise comparisons were used for all post-hoc analyses. A CTA was considered significant if CS preference was significantly lower than NaCl-injected controls. Unconditioned preferences between Wt and FKO mice (Expt 4a) were compared by t-test. For c-Fos induction (Expt 6), genotype (Wt vs. FKO) and treatment (NaCl vs. LiCl) served as the independent factors with mean cell counts per section as the dependent measure for each brain region.

## Acknowledgements

Supported by NIDCD T32-00044 (AK) and NINS R01-50355 (YZ).

